# Headwinds to Understanding Stress Response Physiology: A Systematic Review Reveals Mismatch between Real and Simulated Marine Heatwaves

**DOI:** 10.1101/2024.08.15.608043

**Authors:** Harmony A. Martell, Simon D. Donner

## Abstract

Laboratory experiments have long been used to guide predictions of organismal stress in response to our rapidly changing climate. However, the ability to simulate real world conditions in the laboratory can be a major barrier to prediction accuracy, creating obstacles to efforts informing ecosystem conservation and management. Capitalizing on an extensive experimental literature of coral bleaching physiology, we performed a systematic review of the literature and assembled a database to identify the methods being used to measure coral bleaching in heating experiments and assess how closely heating experiments resembled marine heatwaves (MHWs) on coral reefs. Observations of the maximum photochemical yield of Photosystem II (*F*_V_/*F*_M_), though not a direct measure of bleaching, vastly outnumbered Symbiodiniaceae density and chlorophyll (μg cm^-2^, pg cell^-1^) observations in the available literature, indicating the widespread misuse of *F*_V_/*F*_M_ as a proxy for coral bleaching. Laboratory studies in our database used significantly higher maximum temperatures, degree heating times (∼ 1.7 ×) and heating rates (∼ 7.3 ×), and significantly shorter durations (∼ 1.5 ×), than MHWs on coral reefs. We then asked whether exposure differences between lab and reef altered the relationship between coral bleaching and heating metrics using the example of hormesis, the biphasic dose response wherein low to moderate doses elicit some benefit, while high doses are deleterious. We fit curves on the data both with and without ecologically relevant heating metrics and found hormetic curves in some response variables were altered with the exclusion of exposures that fell outside of the bounds of MHWs on coral reefs. Differences between lab exposures and real-world MHWs were large enough to alter the relationships, indicating a high likelihood of prediction error. We recommend laboratory-based studies of coral bleaching use ecologically relevant exposures to improve our predictions of the coral physiological response to our rapidly warming oceans.

## INTRODUCTION

Bridging the gap between laboratory experiments and the real world is a well-known challenge in science (Holleman et al., 2020; Winkler and Murphy, 1973). In ecology, the goal of laboratory experiments is often to predict how organisms will respond to environmental change. In the Anthropocene, the stakes are high, with predictions focused on whether organisms can keep pace with climate change (Huey et al., 2012; Urban et al., 2024) in order to drive the conservation efforts needed to preserve species and ecosystem function (Ellis, 2019). Experimental extrapolation from the laboratory to the real world has many challenges, including costs, organism collection and husbandry, achieving and maintaining conditions that mimic the real-world environment and experimental durations that enable sufficient prediction. This is further confounded by a lack of consistency across experiments. These and other issues mean reaching any consensus on comparisons to extrapolate to the real world is fraught and difficult, at best.

The field of coral biology is no exception to these extrapolation challenges. There have been several decades of laboratory investigations of warm water coral dysbiosis, colloquially referred to as coral bleaching, and the biggest experimental shortfalls have been identified (McLachlan et al., 2020). To address the laboratory-real world gap, there has been recent movement in the field toward best experimental practices (Grottoli et al., 2021). The consequences of prediction failures in the field of coral ecology are dire, given the well-documented sensitivity of corals to warm temperature extremes (Frieler et al., 2012; Hoegh-Guldberg et al., 2007). While the outlook is grim for coral reefs under business-as-usual warming, there are stress resilience and survival mechanisms that remain unexplored for corals.

The 16^th^ century Swiss alchemist Paracelsus is credited for bringing chemistry and toxicology into the fields of alchemy and early medicine with the phrase *Sola dosis facit venenum*, “Only the dose makes the poison” (1585). This concept posits that exposure to small amounts of poisonous substances may be tolerable, while high amounts are deleterious. This approach is used to define and regulate toxicity in the present day. Dose responses to potential toxins are commonly considered negative, where any amount is detrimental, or neutral where there is a negligible effect. Over the 20^th^ century, the dose response concept has expanded to a variety of scientific disciplines. In the past few decades, thresholds of abiotic environmental variables, such as temperature and light, have being established on organisms in their natural habitat to understand possible ecosystem losses (Erftemeijer and Lewis, 2006; Erftemeijer et al., 2012; Forio et al., 2023; Jensen et al., 2023) and assess climate change risk (Donner, 2009) in the Anthropocene. For example, a common approach used to assess and warn managers of coral bleaching risk involves measures of accumulated heat stress above a local threshold (Liu et al. 2017).

A lesser-known dose response is hormesis. Taken from the Greek word meaning “eagerness”, hormesis is a bimodal response wherein low to moderate doses are stimulating and beneficial rather than neutral, and high doses are harmful. The hormetic model assumes the response to the chemical agent/environmental factor is biphasic, with fitness initially increasing with exposure to the stressor until an asymptote is reached and fitness declines at higher exposures (Costantini et al., 2010). Hormesis has been well studied in toxicology, and is ubiquitous in medicine (Calabrese, 2019; Carelli and Iavicoli, 2016; Mattson, 2008). The concept of hormesis has been applied to understand the influence of abiotic environmental conditions on populations, investigated across a broad variety of ecology disciplines (Costantini et al., 2010), such as agroecology (e.g., Cutler et al., 2022), ecotoxicology (e.g., Butov et al., 2001; Daniels and Allan, 1981; Hansen et al., 1999), and most recently, in assessing organismal climate change resilience (e.g., Costantini et al., 2014; Putnam et al., 2017; Putnam et al., 2020). Hormesis is distinct from acclimation in that, unlike acclimation, hormesis does not presume that the reversible physiological changes induced by the environment will improve future responses: hormesis is only concerned with the prevailing conditions. The hormetic response has been identified as a key driver of phenotypic plasticity which enables organisms to survive extreme events (Costantini, 2014; Costantini, 2019).

Scleractinian corals (hereto forth called corals) are ideal organisms to look for broad evidence of hormesis. Corals live in symbioses with a variety of microorganisms, most notably dinoflagellates from the genus Symbiodiniaceae (Davy et al., 2012; Stat et al., 2006; Weis, 2008). The coral-algal symbiosis makes corals sensitive to environmental perturbations (Weis, 2008). Coral dysbiosis is highly visible: during times of stress, reductions in Symbiodiniaceae density and/or chlorophyll concentration make the coral’s white calcium carbonate skeleton visible through the translucent tissue of the animal (Edmondson, 1928; Glynn, 1984; Glynn, 1996). Corals are thermal specialists with a narrow thermal envelop, leaving them particularly vulnerable to climate change (Angilletta, 2009). Yet, despite a sessile life history, symbiotic corals have persisted since the Triassic period, suggesting they possess a variety of survival mechanisms that enable symbiosis to be maintained (Frankowiak et al., 2021; Stanley and Swart, 1995; Stanley and van de Schootbrugge, 2009). Coral resilience has been linked with hormetic conditioning (Putnam et al., 2020), and hormesis of coral bleaching been demonstrated in the laboratory for a single species (Martell 2022). Hundreds of studies from more than half a century of research have sought to understand coral bleaching in response to elevated temperatures (Davy et al., 2012), particularly in the context of warming (McLachlan et al., 2020).

This long history of coral bleaching studies provides an excellent opportunity to look for broad evidence of thermal hormesis. However, the applicability of laboratory-based results to coral reef management and conservation, whether for hormesis or other physiological phenomena, depends on the gap between the laboratory and the real world. For example, the experimental conditions in such studies must be sufficiently representative of the range of real-world conditions to be instructive for management efforts and interventions aimed at maintaining or enhancing coral reef resilience to climate change. A false positive or false negative finding from laboratory experiments could lead to poor allocation of management resources or maladaptations which reduce coral reef resilience.

This study sought to determine the applicability of laboratory-based warm water coral bleaching experiments to real world conditions, evaluating the evidence with hormesis as a case study. Using a systematic review of the coral bleaching literature, we compared heating metrics from laboratory experiments to MHWs on coral reefs. We then assessed the effect of heating on the coral bleaching response and modeled coral dysbiosis as a function of four heating metrics to detect a hormetic response of the coral holobiont to thermal exposure. We tested four hypotheses. First, we hypothesized that laboratory-based heating metrics would be consistently exaggerated compared to those found on coral reefs during MHWs. Second, we hypothesized a consistently dysbiotic (i.e., negative) effect of heating on coral holobionts, across multiple species. We anticipated that the relationship of coral dysbiosis would be altered by excluding ecologically relevant observations (i.e., removing observations with exposures outside of those occurring on coral reefs during MHWs), pointing to prediction errors when laboratory heating exposures were unrealistic. Thus, we hypothesized at least one hormetic curve would be detected in the coral physiological response to warming, but only when ecologically relevant exposures were applied.

## METHODS

### Systematic Literature Search

We performed a systematic review of the coral bleaching literature to develop a database of time series of temperature and coral bleaching responses from experimental coral bleaching treatments following the procedures of Foo et al. (2021). Exploiting the wealth of coral bleaching response literature, a Web of Science literature search was performed in June 2020 (last date of search: June 19, 2020), using the terms “coral” AND “bleach*” AND “temperature” to identify all relevant studies (Fig 1). The initial search returned 1,584 papers which were then refined by removing papers that were published before 1900, were not written in English, and had not been peer-reviewed (e.g., this excluded conference proceedings). Any titles that indicated the study was not related to coral bleaching were also removed at this stage, resulting in 1,454 papers. A team of three trained reviewers independently screened the abstract of each paper, retaining 359 papers that quantified at least one bleaching response variable (i.e., Symbiodiniaceae density, Chlorophyll, and/or *F*_V_/*F*_M_). The number of papers that quantified each response variable are listed in Table 1. At this stage, all studies were assigned a unique study ID that did not change throughout the process and all 359 papers were screened for the presence of temperature measurements.

**Figure 1.**
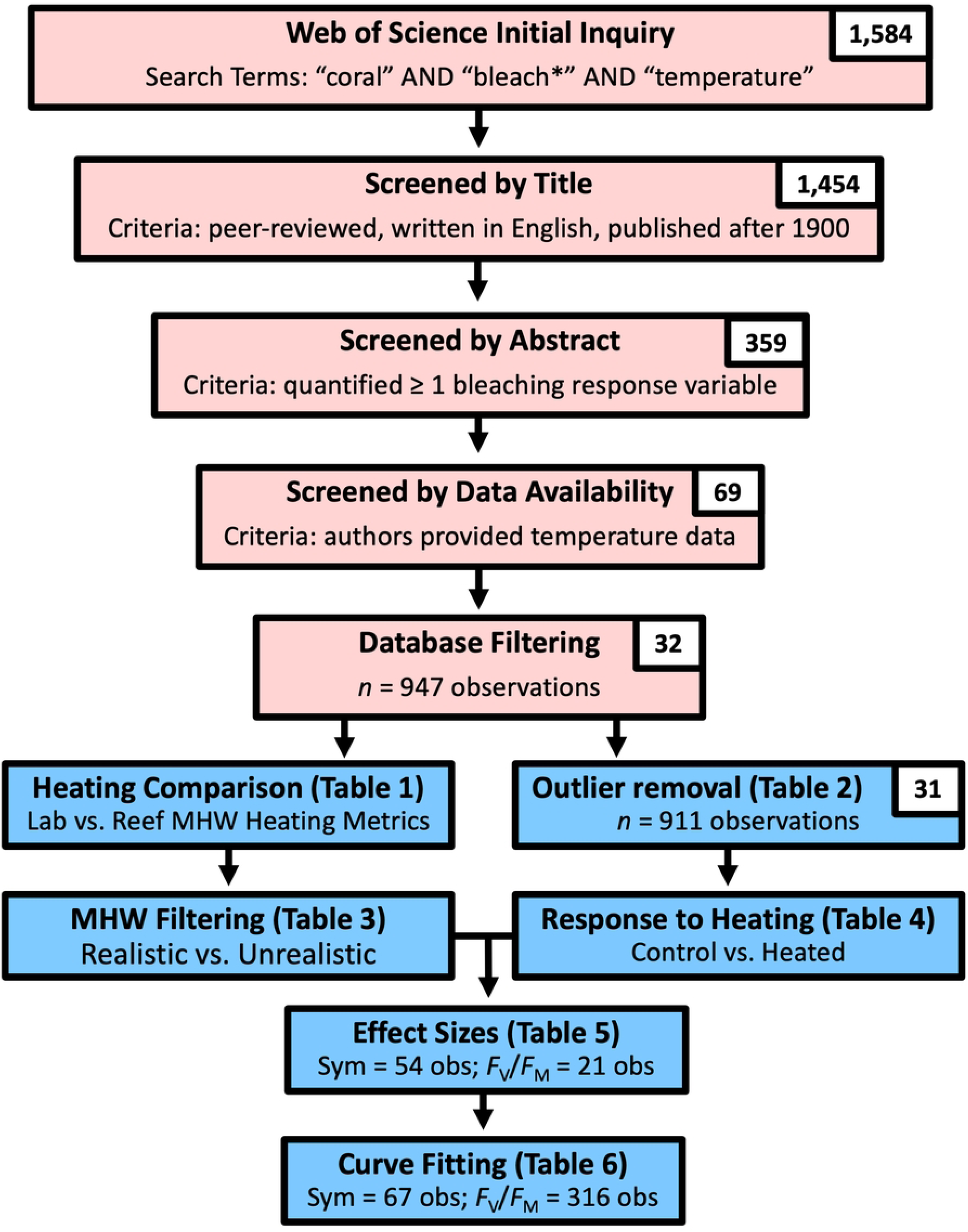
A schematic of the process used in our study of database creation and filtering (pink) and analyses (blue) displaying the number of studies and observations retained at each step.

**Table 1.** The Kruskal Wallis test statistic (𝝌^2^), degrees of freedom (*df*) and *p* value (*p*) for the comparison of heating metrics in the database of laboratory studies from the literature and on coral reef grid cells during marine heatwaves (from Li and Donner, 2022). Significant differences are bolded.

| Heating Metric | $\chi^2$ | $df$ | $p$ |
| --- | --- | --- | --- |
| Maximum Temperature | 366.3 | 1 | **1.18E-81** |
| Heating Duration | 6.8 | 1 | **0.009** |
| Heat Accumulation | 104.52 | 1 | **1.55E-24** |
| Heating Rate | 436.28 | 1 | **6.98E-97** |

### Database Creation

The database was populated by one of four trained reviewers reading through a study thoroughly and then recording qualitative and quantitative information into each of 95 columns, broken into sections including (1) Study Metadata; (2) Experimental Data; (3) Treatments, Variables, and Replicates; (4) Experimental Timeline; (5) Temperature Data; (6) Light Data; (7) Response Variable Data; and (8) Comments. A complete listing of each column, its detailed descriptions and instructions can be found at https://github.com/harmonymartell/Temperature-Coral-Bleaching-Database/commit/bcab1b8b18e9c977b9219f453040d8f7cde6855a.

If the data were not publicly available, the study authors were contacted via email and raw temperature and response variable data was requested using a standard letter and datasheet found at https://github.com/harmonymartell/Temperature-Coral-Bleaching-Database/commit/e9ebfd7ef3c69de8423e54d381cdd2d5cfb6e470. Some studies did not report in situ temperature data. If no in situ temperature data were available from a laboratory study, and there was a sufficient description of the temperature profile reported in the study (e.g., starting temperature, heating rate, heating duration, and maximum temperature at the time the response variable observation was made), we assumed a stable temperature or heating rate and estimated the temperature profile. For a given study, there were *n* number of observations, determined by the mean response variable for a given treatment and sampling time. For example, if Symbiodiniaceae density was quantified at 0, 2 and 5 days in both a control and heated treatment, this would yield a total of six observations, from 2 treatments × 3 timepoints.

Of the 359 studies, 6% provided open access data and additional data was furnished by authors via email contact (12%). For the remaining studies, the authors no longer had the raw data, declined to provide it, or did not respond to the request. A total of 69 articles met our study criteria, resulting in a database of 2,998 observations comprised of a single temperature series and corresponding response variable data from heated or control treatments. The total number of observations across all response variables was higher than the number of observations for any one response variable alone, because some studies measured multiple response variables.

The response variables in this study were physiological observations of coral bleaching metrics (Symbiodiniaceae density, Chlorophyll *a*, Chlorophyll *c_2_* and total Chlorophyll in μg cm^-2^, and Chlorophyll *a* in pg cell^-1^). Mean response variables were quantified for each observation in all studies. The experimental methodology used to quantify response variables was recorded for every observation to ensure comparability across studies (Harrier et al. 2020).

Maximum dark-adapted photochemical yield of photosystem II (*F*_V_/*F*_M_) measurements are a measure of photobiology of the Symbiodiniaceae living in the coral tissues. *F*_V_/*F*_M_ measurements have become ubiquitous in the coral literature because they are measured non-invasively. Therefore, though not a direct measure of coral bleaching, we collected observations of *F*_V_/*F*_M_ if they had corresponding temperature series. As there are several terms used in the coral literature described as *F*_V_/*F*_M_ measurements, to avoid including measurements that were something other than maximum dark-adapted photochemical yield of photosystem II (*sensu* Baker, 2008), we recorded the exact terminology used by the authors to describe the measurement. *F*_V_/*F*_M_ requires measurement of the organism after a period in darkness, referred to as dark-adapting, thus we also recorded whether and the duration of the dark-adapting period for each observation. Additionally, since Symbiodiniaceae have well-documented diurnal and circadian rhythms (Jones and Hoegh-Guldberg, 2001; Sorek et al., 2013), we recorded the time of day the observation was made.

The initial database from the literature collection effort resulted in 2,998 observations of at least one mean response variable and corresponding temperature series made available from the authors, from 69 studies published in 34 different journals from 1996 to 2020 (available at https://github.com/harmonymartell/Temperature-Coral-Bleaching-Database/blob/main/_1_unfilteredDatabase.csv). Observations in the unfiltered database were obtained from experiments performed between 1994 to 2018 with organisms collected or observed at 89 different sites.

For every heated response variable observation, the corresponding maximum temperature, heating duration, heat accumulation, and heating rate were calculated from the raw temperature series of the observation. Calculations and protocols for all response variables and temperature variables are described below.

### Data Processing

All data processing and calculations were performed in R (R Core Team 2020, version 4.0.3). The mean response variable values from all observations were recorded directly from the paper if reported or calculated from the raw data using a custom script. A detailed description of the formatting, processing, and calculation can be found at https://github.com/harmonymartell/Temperature-Coral-Bleaching-Database/blob/main/ResponseVariableDataProcessingProtocol.docx The raw temperature data, consisting of date, time or datetime and temperature from each study were parsed, formatted and processed as follows. First, using the information reported in the study, the temperature data was truncated, leaving only the temperature series that the organism was exposed to, from the start of the experiment to the sampling timepoint that corresponded to the response variable(s) for the given observation. These data were then formatted and the resulting files from heated observations were processed to calculate the heating metrics.

Maximum temperature (°C), heating duration (d), heat accumulation (°C·d) and heating rate (°C d^-1^) for each observation were calculated using a custom script. The maximum temperature was defined as the highest temperature at any point in the temperature series. The heating duration was defined as any time during the temperature series where the temperature was greater than the starting temperature. Heat accumulation was calculated in degree heating days using the time and magnitude above a threshold, typically defined as 1°C above the mean monthly maximum (MMM) temperature (Wyatt et al., 2020). However, due to the complexity in accessing the MMM for each observation based on location data, we used the starting temperature, defined as the first value in the temperature series based on the data provided before any heating began. Thus, all degree heating days were calculated as the heat accumulation above the starting experimental temperature divided by the heating duration. Heating rates were calculated to avoid underestimation, by first truncating all temperature values after the first instance of the maximum temperature and the last minimum temperature that preceded the first instance of the maximum temperature in the series. The duration was calculated as the time between the last minimum temperature value and the first maximum temperature value. Heating rate was then calculated on the parsed temperature series by taking the range and dividing by the corresponding duration.

At this stage, the temperature series of each heated observation was plotted and visually inspected to ensure the temperature series was increasing. Some temperature series from observations that were classified at heated treatments had no change in temperature, precluding the calculation of heating metrics, so they were removed. An exemplary plot of the temperature series can be found in the Supplemental Information (Fig S1) and all temperature series plots for heated observations can be found at https://figshare.com/s/deecce89b3de44ab6eb5.

### Database Filtering

Prior to further analyses, the database was filtered *a priori* to meet our study criteria. All non-coral observations were removed (*n* = 700). Next remaining observations of life stages other than those of adult corals were removed (*n* = 87). Since the intent of our study was solely to identify thermal hormesis, the remaining observations from studies that explicitly altered additional variables (i.e., multiple stressors experiments; *n* = 228) were removed from the database to avoid confounding the thermal stress response. Because few field studies had in situ temperature data and we could not predict the way comparing coarser resolution satellite-derived SST data (i.e., 0.05° x 0.05° deg lat-lon) to in situ data would affect analysis, any remaining observations that were field-based were removed (*n* = 888). The lower limit of Scleractinian coral survival has been reported to be 15°C (Kemp et al., 2016), so remaining observations that were made at temperatures < 15°C (*n* = 6) or exposed corals to ‘decreasing’ or ‘decreasing variable’ temperatures (i.e., cold water bleaching, *n* = 75) were also removed. It is unknown whether facultatively symbiotic corals have the same response to thermal stress as those in obligate mutualism (Holland and Bronstein, 2008; Rivera and Davies, 2021), thus remaining observations with known facultatively symbiotic cnidarian species were removed (*n* = 2 from S0094).

A total of 11 names were used to describe *F*_V_/*F*_M_ observations in our database. *F*_V_/*F*_M_ observations were filtered by removing observations that measured something other than *F*_V_/*F*_M_ (e.g., observations from a single study, S0202, measured Yi and not *F*_V_/*F*_M_), did not report dark-adapting before measuring, did not specify what was measured, or did not report the length of time corals were dark-adapted (*n* = 103).

Observations were examined as a function of ocean province (i.e., Caribbean Sea, Atlanta Ocean, Pacific Ocean, Indian Ocean, or Mediterranean Sea). Nearly all observations were made on tropical corals, thus, remaining observations of corals from the Mediterranean Sea (S0017) were removed (*n* = 4). Database filtering excluded 68 % of the initial 2,998 observations, resulting in a filtered database of 947 observations of at least one response variable with corresponding temperature series from 32 studies. Note that some studies reported multiple response variables for a single observation, while others only had a single response variable. The filtered database was used in downstream exploratory analyses (available at https://github.com/harmonymartell/Temperature-Coral-Bleaching-Database/blob/main/_2_filteredDatabase.csv).

### Marine Heatwave Comparison

The laboratory-based heating metrics from the database developed in this study (*n* = 401) were compared to those of real-world MHWs, using a 2010-2018 warm-season marine heatwaves dataset developed for global 0.05° x 0.05° lat-long coral reef grid cells (*n* = 6,616,000; Li and Donner, 2022). Maximum temperature of MHWs was calculated as the peak hot spot value (°C) plus the climatological maximum monthly mean temperature value (°C) from Li and Donner (2022), using NOAA’s CoralTemp version 3.2 (Skirving et al., 2020). The heating duration (d) was the time the hotspot persisted, the degree heating days were the heat accumulation above the local MMM over the duration, and the heating rates were calculated as the difference in temperature between the hotspot peak and the local MMM divided by the number of days it took to reach the hotspot peak from the beginning of the MHW. Maximum temperature (°C), duration (d), heat accumulation (°C·d) and heating rates (°C d^-1^) from the laboratory-based dataset of this study and Li and Donner (2022) had equivalent units and thus were directly compared. Heating metrics in each treatment group were examined for normality via normal quantile plots and Shapiro-Wilk tests (Shapiro and Wilk, 1965) using *shapiro.test* function and for equal variances using the *var.test* function. *T* tests or Kruskal Wallis tests were performed to test if laboratory experimental designs differed from real world MHWs occurring on coral reefs.

### The Coral Physiological Response to Warming

Observations from the filtered database were parsed by response variable (i.e., Symbiodiniaceae density, Chlorophyll *a* μg cm^-2^, Chlorophyll *a* in pg cell^-1^, total Chlorophyll in μg cm^-2^ and Chlorophyll *c_2_* in μg cm^-2^, and *F*_V_/*F*_M_). The density distributions and boxplots of all variables were examined as a function of treatment (heated vs. control) and examined for possible outliers. The measurement and quantification methods for each response variable were compared between studies to ensure consistency between methods. If a study used an incomparable methodology that produced strongly outlying response variables, observations from that study were removed at this point. The filtered database with no outliers was used to repeat the plotting, and for downstream analyses (Fig 1, Blue Boxes).

All functions used in exploratory hypothesis testing were from the *stats* package or base R. Response variables in each treatment group were examined for normality via normal quantile plots and Shapiro-Wilk tests (Shapiro and Wilk, 1965) using *shapiro.test* function, and for equal variances using the *var.test* function. Kolmogorov-Smirnoff tests (Chakravart, Laha, and Roy, 1967) were performed to assess whether samples from control and heated treatments were drawn from the same distribution using the *ks.test* function. Wilcoxon rank sum tests with continuity corrections (Bauer, 1972) were used to determine whether control treatment medians were significantly greater than heated treatment medians via the *wilcox.test* function. Significant differences between treatments were identified by either ANOVA when the data met assumptions, using the *aov* function, or via a Kruskal Wallis test (Kruskal and Wallis, 1952) when the data did not meet assumptions, using the *kruskal.test* function. To ensure compatibility between hypothesis tests, a subset of 201 *F*_V_/*F*_M_ observations from 13 studies that also made observations of Symbiodiniaceae density were taken and the same statistics were performed.

The percentage of observations that fell within what was observed in the MHW dataset were then calculated to ascertain ecological relevance, that is, how many observations were from experiments that used “realistic” (i.e., within the range observed in real-world MHWs on coral reefs, henceforth referred to as ecologically relevant) vs. “unrealistic” heating metrics. To create the ecologically relevant subset, the database observations were filtered by the maximal or minimal MHW values from coral reef grid cells to retain observations. The combination of both maximum temperature and duration was used to filter observations for the ecologically relevant subset; heat accumulation measured in degree heating times was not used because identical values can represent different exposures (high temperature increases over shorter durations vs. moderate temperature increases over longer durations).

### The Effect of Heating on Response Variables

Standard mean differences (SMD), or Hedge’s G, were calculated for each coral physiological response variable to determine the proportion of change between heated and control treatments in response to heating (*esc* package, Lüdecke 2018) according to (2021). Effect sizes (i.e., SMD), their respective weights based on sample sizes, and the sample variances of each effect size were calculated for each experiment. In studies that included multiple species, each species was considered as a separate experiment since distinct comparisons were made between heated and control treatments for each species. In the case where a response variable lacked sufficient sample size to determine an effect, these response variables were removed from downstream analysis.

### Hormetic Curve Fitting

The heated observations from the remaining response variables (i.e., those for which a clear negative effect could be determined) and their corresponding heating metrics were taken from the database. Using the response variable as measures of symbiotic association strength (i.e., fitness), we modeled the heated physiological response as a function of each heating metric, to detect a hormetic dose response to warm water exposure. We fit a 2^nd^ degree polynomial on the relation of each response variable (e.g., Symbiodiniaceae density, Chlorophyll *a*, or *F*_V_/*F*_M_) as a function of each heating metric (e.g., maximum temperature, heating duration, heat accumulation, or heating rate).

Nonlinear mixed effects models were constructed to examine response variables as a function of heating metric (fixed effect; *lme4* package, Bates et al. 2011). To account for the inherent variation between observations (e.g., oceanic provinces, coral species, local adaptation, and acclimatization, etc.), models were conducted with study ID as the random effect. Models were assessed for normality and heteroscedasticity. Model singularity was recorded, as it can indicate too few random effects (i.e., too few studies). Predictor variables (i.e., heating metrics) were log transformed when their scale was too broad. An ANOVA was performed to determine whether the slopes of each model (𝑎𝑥^2^ and 𝑏𝑥) were significantly different from zero at α = 0.05. If there were significant slopes for both model coefficients, with a positive 𝑎𝑥^2^ coefficient and negative 𝑏𝑥 coefficient, the relation was identified as hormetic. The marginal and conditional *R*^2^ were calculated for the final models to estimate the variance explained by heating metric, and heating metric plus study ID, respectively.

Fitting was then repeated using only the ecologically relevant subset of observations to test whether the use of unrealistic heating metrics altered the relationship of the physiological response to warming. Model coefficients from the complete and ecologically relevant fits were compared to determine whether the relationships differed. All scripts to complete the analysis are available at https://github.com/harmonymartell/Temperature-Coral-Bleaching-Database/tree/main.

## RESULTS

### Marine Heatwave Comparison

All heating metrics in the laboratory were significantly different from those during MHWs on coral reefs (Fig 2, Table 1). Maximum temperatures, degree heating days, and heating rates used in laboratory experiments (*n* = 401) were significantly greater, and laboratory heating durations were significantly shorter, than those made during MHWs (*n* = 5,629,990). The median maximum temperature employed in laboratory observations (μ_1/2_ = 32.3 ± 0.2 °C) was significantly greater than the median MHW maximum temperature (μ_1/2_ = 30.4 ± 0.0 °C). The median laboratory heating duration was 21.5 ± 5.8 days compared to the significantly longer median MHW duration of 30.0 ± 0.1 days. Accumulated heating was nearly two fold higher in laboratory values compared to accumulated heating during MHWs, with median laboratory exposures of 39.7 ± 19.0 °C·d and MHW median degree heating days at 24.0 ± 0.01 °C·d. Median laboratory heating rates (μ_1/2_ = 0.51 ± 0.29 °C d^-1^) were more than seven times greater than those observed in coral grid cells during MHWs (μ_1/2_ = 0.07 ± 0.00 °C d^-1^). The maximum heating rate observed during a MHW on a coral reef grid cell was 1.52 °C d^-1^, thus this value was set at the threshold for downstream analyses.

**Figure 2.**
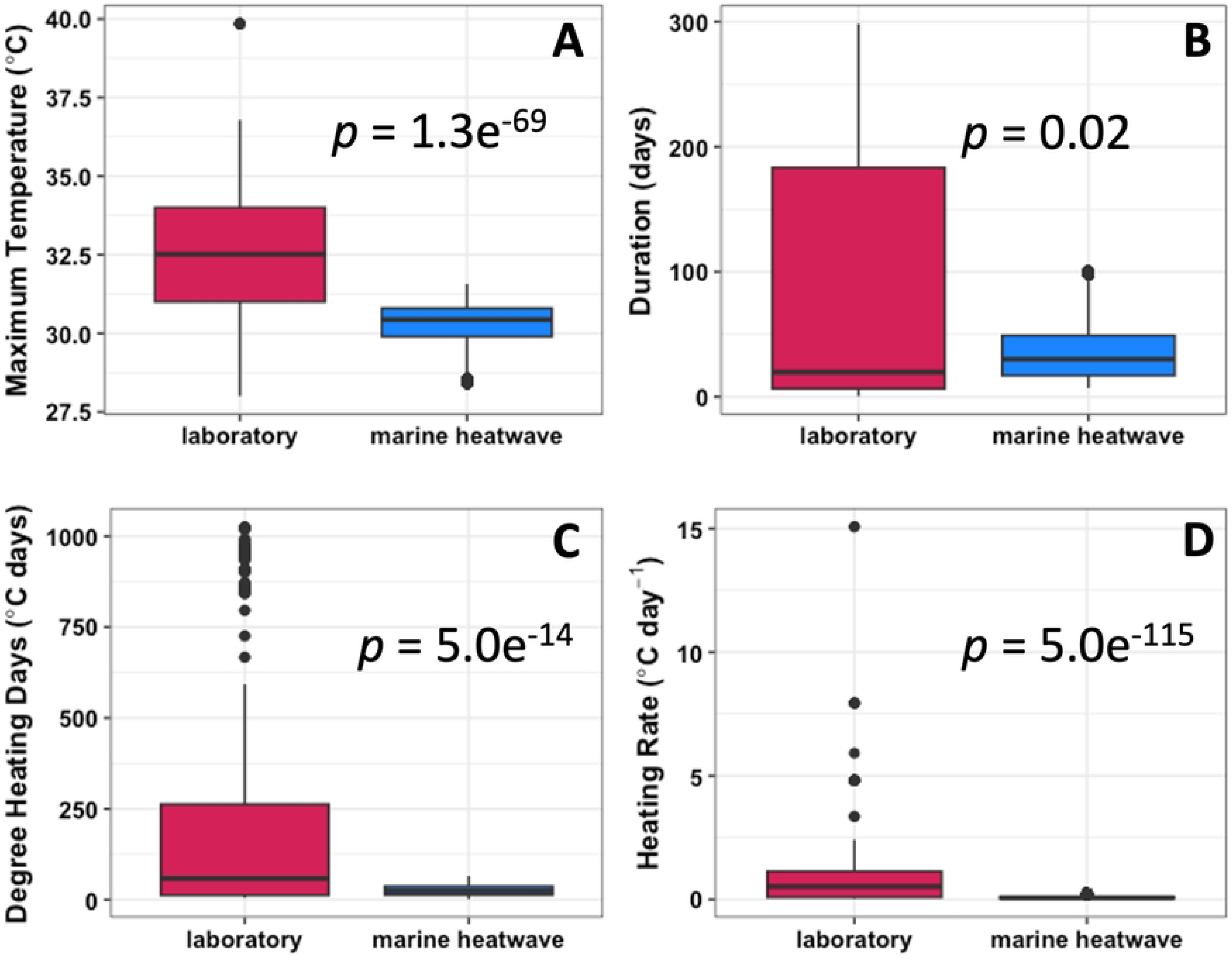
The comparison of the heating metrics (A) maximum temperature, °C, (B) heating duration, d, (C) heat accumulation, °C·d, and (D) heating rate, °C d^-1^ from database studies (laboratory, dark pink) and coral reef grids cells during marine heatwaves (marine heatwaves, blue). Data shown are the 95%ile.

### Database Response Variables

*F*_V_/*F*_M_ was the most frequently reported response variable (*n* = 778), with Symbiodiniaceae density being the next highest (*n* = 245), followed by Chlorophyll *a* (pg cell^-1^, including reported values and values calculated by this study, *n* = 122), and Chlorophyll *a* (μg cm^-2^, *n* = 67; Table 2). Total Chlorophyll (*n* = 14 observations from 2 studies) and Chlorophyll *c_2_* (*n* = 13 observations from 4 studies) had the least number of observations (Table 2). Exploratory analyses revealed a single study (S0093) had outlying values for both Symbiodiniaceae density and Chlorophyll *a* per cell that were greater than all other database entries, due to a difference in the quantification methods used to obtain these values. Because no correction could be applied, the study was removed from the database (*n* = 36 observations).

**Table 2.** The numbers of observations used at each analysis step. The number of observations by treatment (*n,* control, heated) from the database used for exploratory analyses, to determine effect sizes, and in hormetic curve fitting. Hormetic fitting was performed using the full dataset (Complete) and the dataset that was filtered for ecologically relevant heating metrics (Filtered). -- indicates the analysis was not performed. Note that the number of observations from fitting before filtering (Complete) are less than heated observations in the exploratory analysis. This is because exploratory analysis was undertaken on all samples in the database, even if temperature data were not available for them. Additionally, some temperature series classified as heated had no change in temperature, which precluded calculation of heating metrics.

| ***n* Observations** | **Exploratory Analyses** |  | **Effect Size Calculations** | **Hormetic Fitting** |  |
| --- | --- | --- | --- | --- | --- |
| Response Variable | Control | Heated | Control-Heated Pairs | Complete | Filtered |
| Symbiodiniaceae Density (cells cm<sup>-2</sup>) | 121 | 88 | 12 | 67 | 34 |
| Chlorophyll *a* (μg cm<sup>-2</sup>) | 41 | 26 | 0 | 15 | 13 |
| Chlorophyll *a* (pg cell<sup>-1</sup>) | 55 | 31 | 1 | 19 | 13 |
| Chlorophyll *c*<sub>2</sub> (μg cm<sup>-2</sup>) | 7 | 6 | 0 | -- | -- |
| Total Chlorophyll (μg cm<sup>-2</sup>) | 10 | 4 | 0 | -- | -- |
| *F<sub>V</sub>*/*F<sub>M</sub>* (all data) | 339 | 336 | 9 | 316 | 230 |
| *F<sub>V</sub>*/*F<sub>M</sub>* (subset) | 116 | 118 | -- | -- | -- |

Only 51% of the heated observations of Symbiodiniaceae were made using ecologically relevant exposures (Table 3). A higher fraction of heated observations of Chlorophyll *a* in μg cm^-2^ and pg cell^-1^ were within MHW exposure values (87 % and 68%, respectively, Table 3), though there were fewer Chlorophyll *a* observations overall. Roughly two-thirds of heated *F*_V_/*F*_M_ observations used realistic heating metrics. Observation counts for each response variable at each filtering step can be found in Table 3.

**Table 3.** The number of observations that met the filtering criteria used to assess the proportion of laboratory observations that used ecologically relevant exposures, based on SST observations made on coral reefs during marine heatwaves from 2010-2018. The number of observations that fell within maximum temperature (Max. Temp.) values on marine heatwaves, the duration (Duration), both maximum temperature and duration (Temp. Duration), and heating rates (Rate), as well as the remaining amount that met all ecologically relevant heating criteria (All), the starting number of observations (Total) and the percentage of observations that were ecologically relevant (% Realistic) for each Response Variable in the database.

| **Response Variable** | **Max. Temp.** | **Duration** | **Temp. Duration** | **Rate** | **All** | **Total** | **% Realistic** |
| --- | --- | --- | --- | --- | --- | --- | --- |
| Symbiodiniaceae Density | 66 | 49 | 48 | 40 | 34 | 67 | 51% |
| Chlorophyll *a* ($\mu\text{g cm}^{-2}$) | 15 | 15 | 15 | 13 | 13 | 15 | 87% |
| Chlorophyll *a* ($\text{pg cell}^{-1}$) | 19 | 19 | 19 | 13 | 13 | 19 | 68% |
| $F_V/F_M$ | 291 | 287 | 262 | 272 | 230 | 316 | 73% |

### The Coral Physiological Response to Warming

The density distributions of heated versus control Symbiodiniaceae density and *F*_V_/*F*_M_ differed in both the distribution shape and distribution median (Table 4, Figs S2A, S3A). Mean heated treatment values were consistently lower than the controls for Symbiodiniaceae density and *F*_V_/*F*_M_ (Table 4, Figs S2B, S3B). This trend followed for the subsetted *F*_V_/*F*_M_ distributions and treatment differences as well (Table 4, Fig S4). There was a marginal difference in median distributions between treatments of Chlorophyll *a* (μg cm^-2^), though the treatment distribution shapes and medians did not differ (Table 4, Fig S5). Chlorophyll a (pg cell^-1^), Chlorophyll *c_2_*, and total Chlorophyll did not differ between treatments (Table 4, Figs S6-8). Median ± 1SE response variables by treatment and all test statistics are reported in Table 4.

**Table 4.**
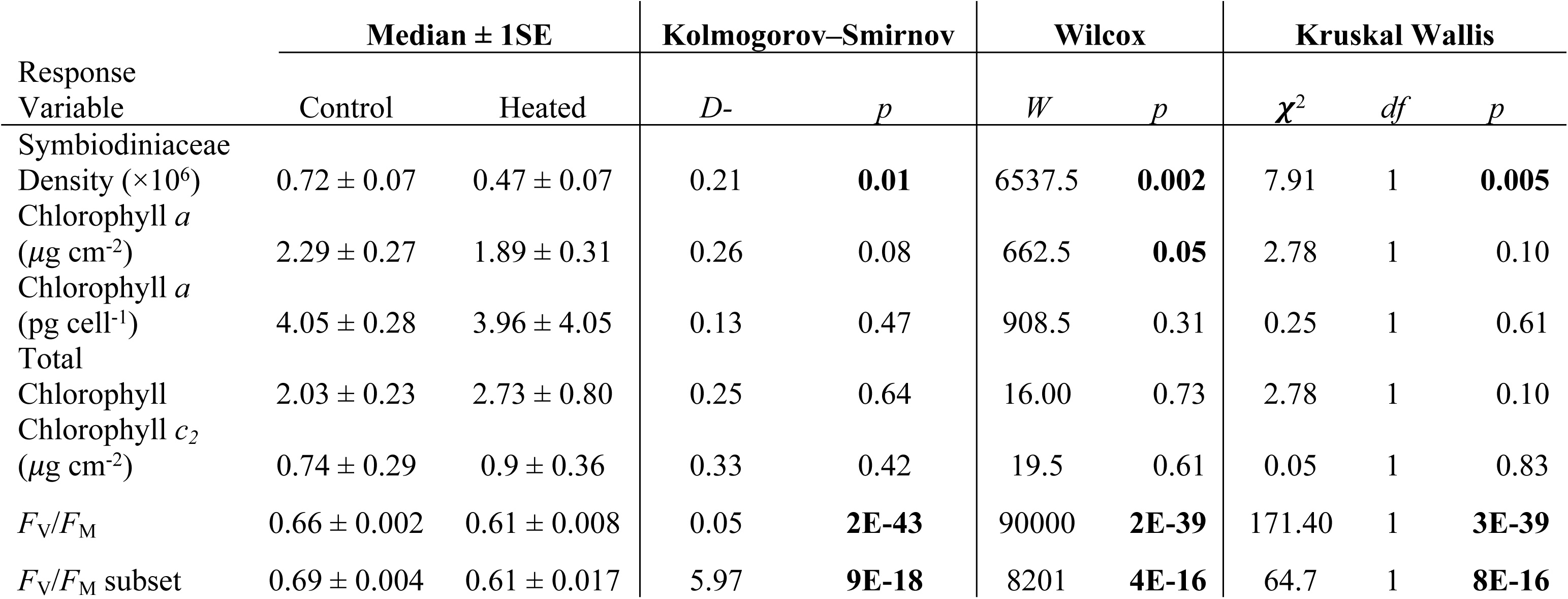
The median ± 1SE treatment (control and heated) values for coral physiological response to warming exposures and corresponding Kolmogorov-Smirnov, Wilcox and Kruskal Wallis test statistics and *p* values for the comparison of treatments. Significant values are bolded.

| Response Variable | Median ± 1SE | | Kolmogorov-Smirnov | | Wilcoxon | | Kruskal Wallis | | |
| --- | --- | --- | --- | --- | --- | --- | --- | --- | --- |
| | Control | Heated | $D$- | $p$ | $W$ | $p$ | $\chi^2$ | $df$ | $p$ |
| Symbiodiniaceae Density ($\times 10^6$) | 0.72 ± 0.07 | 0.47 ± 0.07 | 0.21 | **0.01** | 6537.5 | **0.002** | 7.91 | 1 | **0.005** |
| Chlorophyll $a$ ($\mu\text{g cm}^{-2}$) | 2.29 ± 0.27 | 1.89 ± 0.31 | 0.26 | 0.08 | 662.5 | **0.05** | 2.78 | 1 | 0.10 |
| Chlorophyll $a$ (pg cell$^{-1}$) | 4.05 ± 0.28 | 3.96 ± 4.05 | 0.13 | 0.47 | 908.5 | 0.31 | 0.25 | 1 | 0.61 |
| Total Chlorophyll | 2.03 ± 0.23 | 2.73 ± 0.80 | 0.25 | 0.64 | 16.00 | 0.73 | 2.78 | 1 | 0.10 |
| Chlorophyll $c_2$ ($\mu\text{g cm}^{-2}$) | 0.74 ± 0.29 | 0.9 ± 0.36 | 0.33 | 0.42 | 19.5 | 0.61 | 0.05 | 1 | 0.83 |
| $F_v/F_M$ | 0.66 ± 0.002 | 0.61 ± 0.008 | 0.05 | **2E-43** | 90000 | **2E-39** | 171.40 | 1 | **3E-39** |
| $F_v/F_M$ subset | 0.69 ± 0.004 | 0.61 ± 0.017 | 5.97 | **9E-18** | 8201 | **4E-16** | 64.7 | 1 | **8E-16** |

### Effect Size Calculations

Effect size calculations require paired heated and control observations from the same experiment (in this case, observations from the same study and species), and thus were possible only on a subset of the data (Table 3). Chlorophyll *a* (μg cm^-2^), Chlorophyll *c_2_*, and total Chlorophyll lacked sufficient paired observations to calculate effect sizes. Chlorophyll *a* (pg cell^-1^) had only one paired observation thus the effect size was calculated (y hat = 0.67), but conclusions could not be drawn about the relationship to any heating metric. Symbiodiniaceae density and *F*_V_/*F*_M_ yielded 12 and 9 experiments, respectively, enabling calculation of the effect of heating on those response variables (Table 5). Heating had a negative effect on the nine out of 12 experiments of Symbiodiniaceae density (Table 5A), but only four of twelve outcomes were significant (i.e., an effect is significant only when the confidence intervals do not cross 0). Heating had a negative effect on *F*_V_/*F*_M_ for all nine experiments but only four of the nine outcomes were significant (Table 5B).

**Table 5.** The effect of heating on Symbiodiniaceae density (A) and *F*_V_/*F*_M_ (B) for each study & species combination from the database, including the Effect Size (Hedge’s G), Weight, Sample Size (i.e., number of paired control-heat observations; *n*), standard error (SE), Variance (Var), Lower 95 % Confidence Interval (Lower CI) and Upper 95% Confidence Interval (Upper CI). Confidence Intervals that did not cross zero are bolded.

| Study ID | Species | Effect Size | Weight | $n$ | SE | Var | Lower CI | Upper CI |
| --- | --- | --- | --- | --- | --- | --- | --- | --- |
| S0021 | *Acropora tenuis* | -2.03 | 1.19 | 8 | 0.92 | 0.84 | **-3.83** | **-0.24** |
| S0021 | *Favites colemani* | 0.08 | 2.00 | 8 | 0.71 | 0.50 | -1.31 | 1.47 |
| S0021 | *Montipora digitata* | -1.67 | 1.37 | 8 | 0.86 | 0.73 | **-3.35** | **0.00** |
| S0021 | *Seriatopora caliendrum* | -1.85 | 1.28 | 8 | 0.88 | 0.78 | **-3.58** | **-0.11** |
| S0033 | *Acropora millepora* | -0.67 | 1.38 | 6 | 0.85 | 0.73 | -2.34 | 1.00 |
| S0033 | *Galaxea acrhelia* | -0.59 | 1.41 | 6 | 0.84 | 0.71 | -2.24 | 1.07 |
| S0033 | *Galaxea astreata* | -0.17 | 1.49 | 6 | 0.82 | 0.67 | -1.78 | 1.43 |
| S0033 | *Goniopora columna* | -0.12 | 1.50 | 6 | 0.82 | 0.67 | -1.72 | 1.48 |
| S0033 | *Montipora aequituberculata* | 0.70 | 1.37 | 6 | 0.85 | 0.73 | -0.97 | 2.38 |
| S0033 | *Porites lobata* | -1.39 | 1.09 | 6 | 0.96 | 0.92 | -3.26 | 0.49 |
| S0277 | *Stylophora pistillata* | -1.19 | 4.57 | 24 | 0.47 | 0.22 | **-2.11** | **-0.27** |
| S0382 | *Acropora formosa* | 0.02 | 1.00 | 4 | 1.00 | 1.00 | -1.94 | 1.98 |

B.
| Study ID | Species | Effect Size | Weight | $n$ | SE | Var | Lower CI | Upper CI |
| --- | --- | --- | --- | --- | --- | --- | --- | --- |
| S0012 | *Montipora digitata* | -0.56 | 3.77 | 16 | 0.51 | 0.27 | -1.57 | 0.45 |
| S0021 | *Acropora tenuis* | -1.80 | 1.30 | 8 | 0.88 | 0.77 | **-3.51** | **-0.08** |
| S0021 | *Favites colemani* | -0.54 | 1.91 | 8 | 0.72 | 0.52 | -1.96 | 0.88 |
| S0021 | *Montipora digitata* | -1.52 | 1.45 | 8 | 0.83 | 0.69 | -3.14 | 0.11 |
| S0021 | *Seriatopora caliendrum* | -1.18 | 1.63 | 8 | 0.78 | 0.61 | -2.72 | 0.36 |
| S0054 | *Acropora millepora* | -0.96 | 4.44 | 20 | 0.47 | 0.23 | **-1.89** | **-0.03** |
| S0106 | *Acropora aspera* | -0.63 | 10.84 | 60 | 0.30 | 0.09 | **-1.22** | **-0.03** |
| S0128 | *Porites compressa* | -1.26 | 2.85 | 14 | 0.59 | 0.35 | **-2.42** | **-0.10** |
| S0382 | *Acropora formosa* | -0.57 | 0.89 | 4 | 1.06 | 1.13 | -2.65 | 1.51 |

Traditional effect size calculation techniques required paired observations of a given response variable for both control and heated treatments, requiring the removal of even more observations from the database, reducing the number of available observation to include in analyses. After the calculation of effect sizes, the experiments in our database lacked evenness across heating metrics to examine the effect of each metric on the outcome (Figs. S9, S10) before exclusion of observations with heating metrics that were not ecologically relevant. Further, when observations that were not ecologically relevant were removed, in all cases, there were insufficient samples spanning the range of predictor variables (i.e., heating metrics) to confidently construct a model.

Exposure to warming is widely known to result in the reduction of Symbiodiniaceae density and *F*_V_/*F*_M_ in corals (Brown, 1997; Douglas, 2003) and observations from our database were in support of this, with consistently reduced density distributions and distribution medians in the heated corals relative to controls (Figs 2-8A). By definition, hormesis occurs with sublethal, rather than marked stress (Berry and López-Martínez, 2020; Tang et al., 2019). There was a significant negative effect of heating in 33% and 44 % of Symbiodiniaceae density and *F*_V_/*F*_M_ experiments, respectively. The application of traditional meta-regression techniques would leave too few observations to compare the influence of ecological relevance of exposures on the relationship of any response variable vs. heating metric. For all these reasons, we instead performed curve fitting on heated observations of each response variable against each heating metric, for response variables with sufficient sample sizes to determine a negative effect (i.e., Symbiodiniaceae density and *F*_V_/*F*_M_).

### Hormetic Curve Fitting

Most fits of the unfiltered, complete dataset on response variables were not hormetic using our *a priori* criteria, and yielded coefficients that were not significantly different from zero at α = 0.05, with low *R*^2^ values (Table 6, Figs 3, 4). The relation of unfiltered *F*_V_/*F*_M_ with heat accumulation and heating rate had coefficients that were significantly different from zero, though these relationships were the inverse of a hormetic response (i.e., a negative *ax^2^* slope and a positive *bx* slope, resulting in a U shape; Figs 4B, 4D; Table 6). Heated observations of all Chlorophyll data lacked sufficient observations and number of studies to perform fitting, thus they were excluded (Table 2).

**Figure 3.**
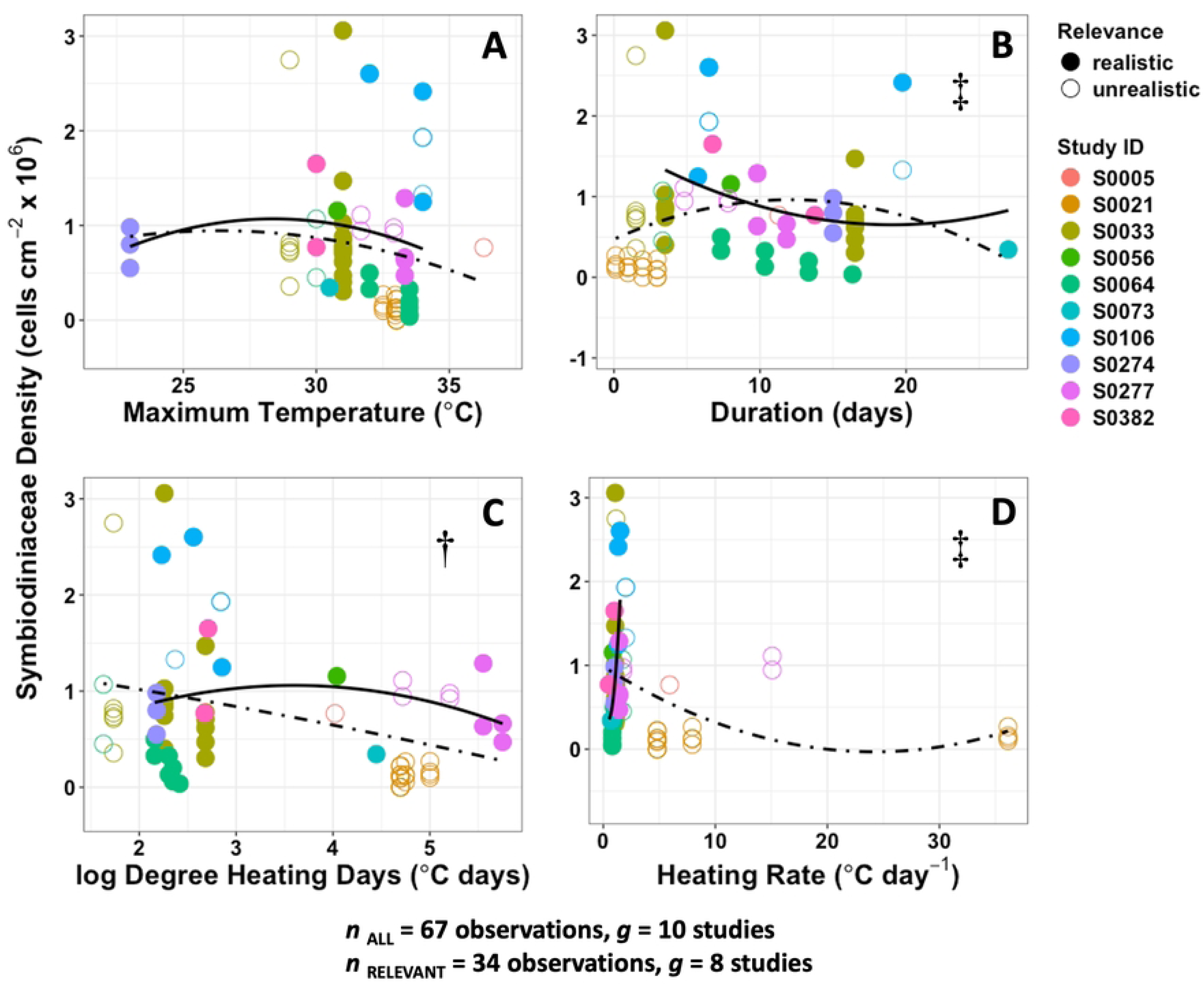
Hormetic fits of heated responses of Symbiodiniaceae density against (A) maximum temperature, (B) duration, (C) heating duration, and (D) heating rate. Dot-dashed lines are fits of complete dataset, while solid lines are fits of the ecologically relevant subset (i.e., those within the limits of exposures observed on coral reefs during MHWs). Solid circles are ecologically relevant observations, and open circles are observations from outside the limits of MHW exposures. † indicates a change in one or more slope directions, * indicates ≥ 2-fold increase in one or more slope magnitudes, ‡ indicates a change in both direction and magnitude.

**Figure 4.**
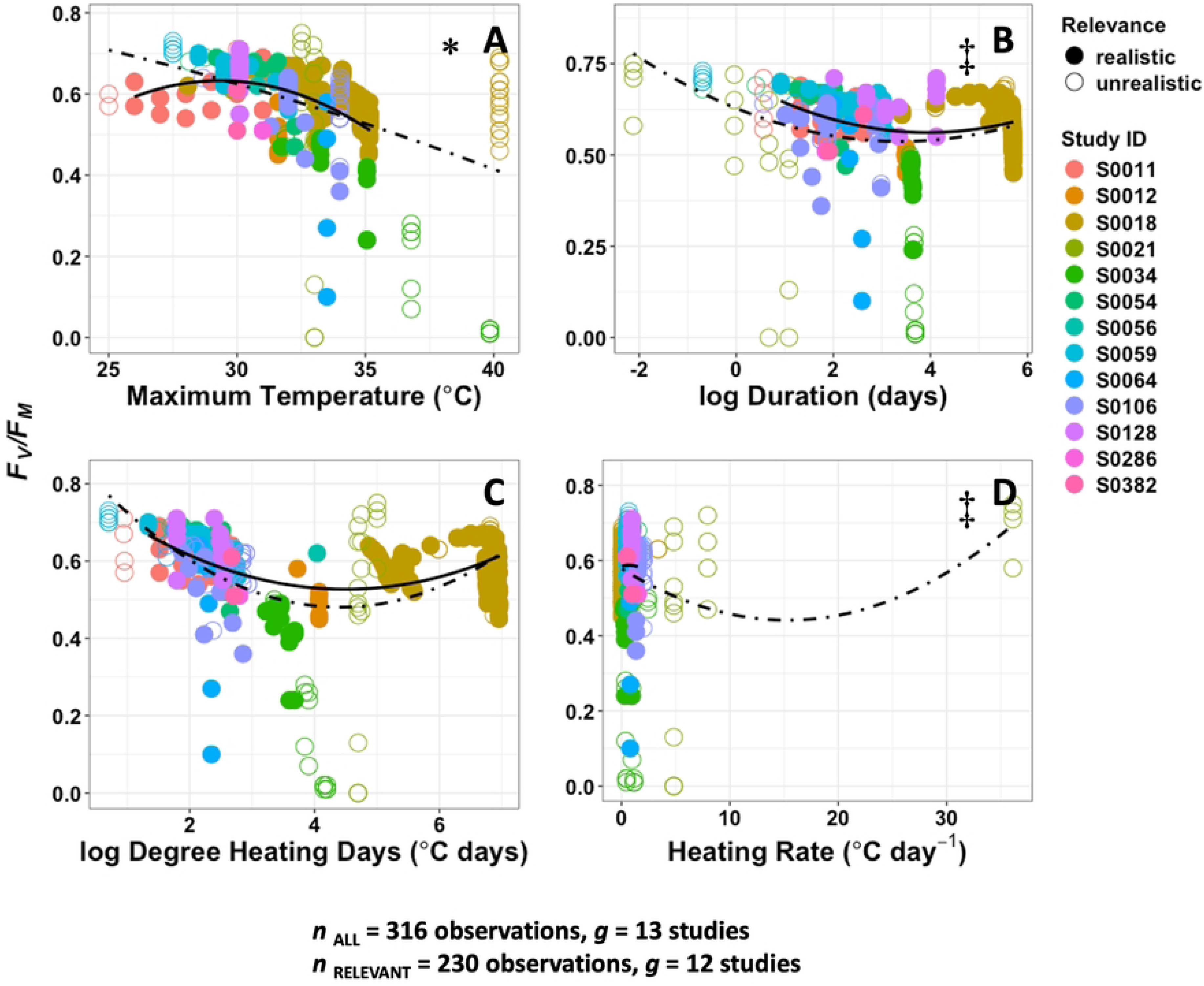
Hormetic fits of heated responses of *F*_V_/*F*_M_ against (A) maximum temperature, (B) duration, (C) heating duration, and (D) heating rate. Dot-dashed lines are fits of complete dataset, while solid lines are fits of the ecologically relevant subset (i.e., those within the limits of exposures observed on coral reefs during MHWs). Solid circles are ecologically relevant observations, and open circles are observations from outside the limits of MHW exposures. † indicates a change in one or more slope directions, * indicates ≥ 2-fold increase in one or more slope magnitudes, ‡ indicates a change in both direction and magnitude.

**Table 6.**
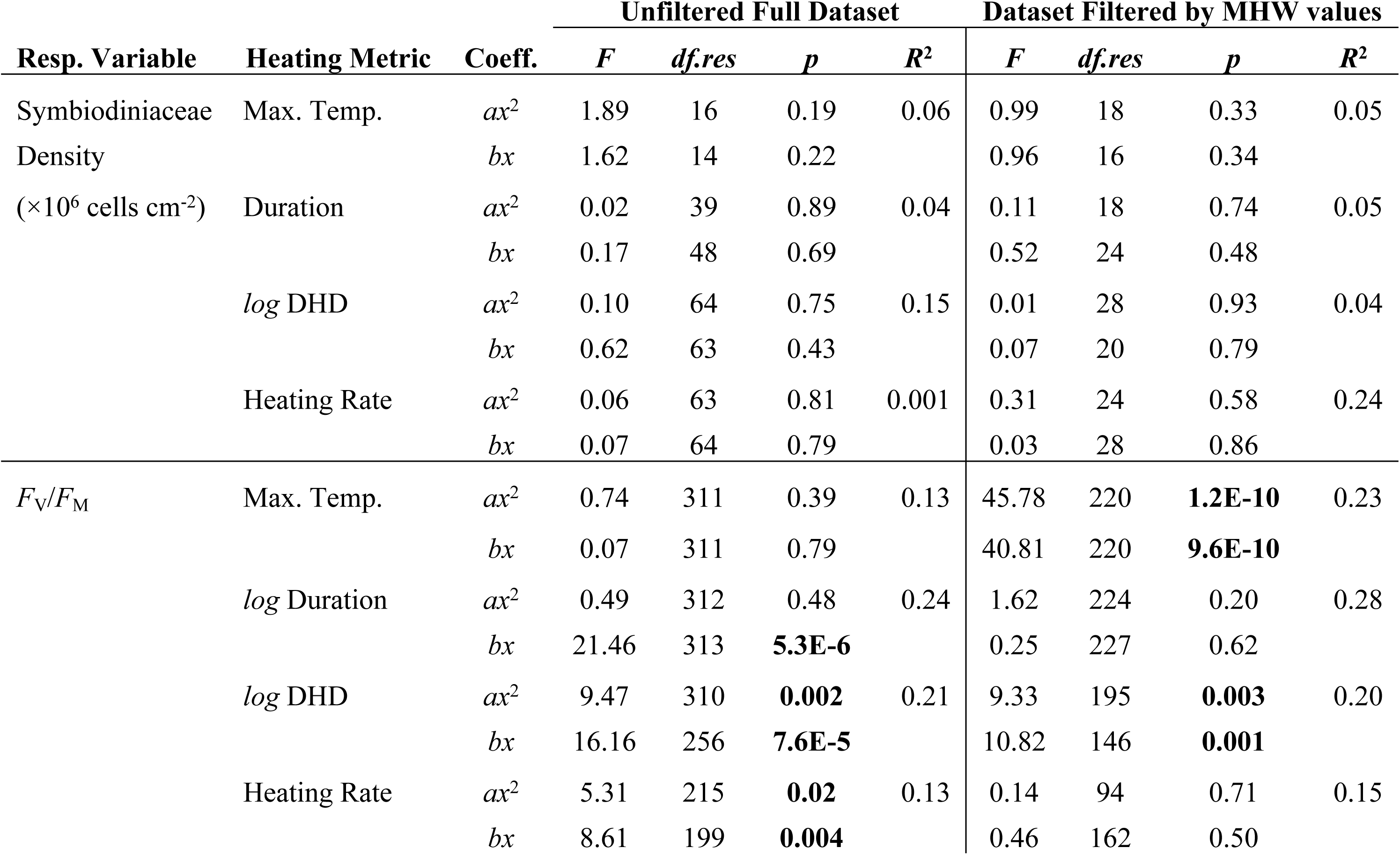
The *F*-statistic, residual degrees of freedom (*df.res*), *p* values (*p*) and *R*^2^ for the hormetic fits of each response variable (Resp. Variable) on each Heating Metric. All models had degrees of freedom = 1. Statistics are reported for both model coefficients (Coeff.) in each fit. Significant values are bolded.

In the ecologically relevant subset, no significant hormesis of Symbiodiniaceae density was detected with any heating metric (Fig 3, Table 6). The fits of the *F*_V_/*F*_M_ data yielded hormetic relationships (i.e., both 𝑎𝑥^2^and ―𝑏𝑥 slopes were significant) with maximum temperature and heat accumulation, though the models poorly explained the variance in the data (*R*^2^ < 30%, Table 6, Figs 4A, C). Of note, *F*_V_/*F*_M_ data had a much larger sample size than Symbiodiniaceae density for fitting (Table 2, Figs 3, 4).

The removal of observations that were made using exposures that were not ecologically relevant resulted changes in the significance of model coefficients (Table 6). For example, the coefficients for the relations of *F*_V_/*F*_M_ vs. heating rate were no longer significant after removing observations that were not ecologically relevant. Additionally, changes in both model coefficient magnitudes (i.e., fold change) and directions (i.e., a change in slope from negative to positive; Table 7, Figs 3, 4) were identified in three out of four models for both Symbiodiniaceae density and *F*_V_/*F*_M_, with at least a 2x change in slope magnitude for one or more coefficients, change in slope direction, or both (Table 7; Figs 3, 4). The relation of Symbiodiniaceae density with The removal of observations that were not ecologically relevant altered the relationships between coral physiological response variables and warming, indicating these predictions likely had a large amount of error.

**Table 7.** The hormetic model coefficients ± 1SE of fits using the complete dataset (All) and only observations that used ecologically relevant heating metrics (Relevant) for Symbiodiniaceae density and *F*_V_/*F*_M_ as a function of heating metrics, along with the change in magnitude (Fold Change), and whether they changed direction. Changes in magnitude > 1 are bolded.

| Resp. Var. | Heating Metric | Coeff. | All | Relevant | Fold Change | Direction |
| --- | --- | --- | --- | --- | --- | --- |
| Symbiodiniaceae<br>Density<br>($\times 10^6$ cells $\text{cm}^{-2}$) | Max. Temp. | $c$ | $-11.38 \pm 10.31$ | $-13.91 \pm 14.64$ | 1 | |
| | | $ax^2$ | $-0.02 \pm 0.01$ | $-0.02 \pm 0.02$ | 1 | |
| | | $bx$ | $0.90 \pm 0.69$ | $1.07 \pm 1.02$ | 1 | |
| | Duration | $c$ | $1.10 \pm 0.27$ | $1.45 \pm 0.45$ | 1 | |
| | | $ax^2$ | $-0.0003 \pm 0.00$ | $0.001 \pm 0.00$ | **-3** | X |
| | | $bx$ | $-0.02 \pm 0.05$ | $-0.06 \pm 0.08$ | **3** | |
| | $\log$ DHD | $c$ | $1.89 \pm 0.83$ | $3.46 \pm 2.19$ | **2** | |
| | | $ax^2$ | $0.02 \pm 0.07$ | $0.15 \pm 0.17$ | **8** | |
| | | $bx$ | $-0.39 \pm 0.48$ | $-1.31 \pm 1.29$ | **3** | |
| | Heating Rate | $c$ | $0.83 \pm 0.21$ | $0.44 \pm 0.90$ | 1 | |
| | | $ax^2$ | $-0.0003 \pm 0.00$ | $1.11 \pm 1.83$ | **-3700** | X |
| | | $bx$ | $0.01 \pm 0.04$ | $-0.73 \pm 3.78$ | **-73** | X |
| $F_V/F_M$ | Max. Temp. | $c$ | $0.67 \pm 0.49$ | $-4.18 \pm 0.79$ | **-6** | X |
| | | $ax^2$ | $-0.0004 \pm 0.00$ | $-0.01 \pm 0.00$ | **25** | |
| | | $bx$ | $0.01 \pm 0.03$ | $0.32 \pm 0.05$ | **32** | |
| | $\log$ Duration | $c$ | $0.66 \pm 0.04$ | $0.63 \pm 0.05$ | 1 | |
| | | $ax^2$ | $0.001 \pm 0.00$ | $-0.004 \pm 0.00$ | **-4** | X |
| | | $bx$ | $-0.05 \pm 0.01$ | $-0.01 \pm 0.02$ | 0 | |
| | $\log$ DHD | $c$ | $0.78 \pm 0.06$ | $0.77 \pm 0.06$ | 1 | |
| | | $ax^2$ | $0.01 \pm 0.00$ | $0.01 \pm 0.00$ | 1 | |
| | | $bx$ | $-0.1 \pm 0.02$ | $-0.09 \pm 0.03$ | 1 | |
| | Heating Rate | $c$ | $0.51 \pm 0.04$ | $0.50 \pm 0.04$ | 1 | |
| | | $ax^2$ | $-0.001 \pm 0.00$ | $0.03 \pm 0.06$ | **-30** | X |
| | | $bx$ | $0.04 \pm 0.01$ | $0.06 \pm 0.09$ | **2** | |

## DISCUSSION

### Laboratory Exposures in the Context of Marine Heatwaves on Coral Reefs

We found significantly higher maximum temperatures and shorter durations used in the laboratory than those of MHWs occurring on coral reefs from 2010-2018. This resulted in significantly higher heat accumulation for corals in the laboratory, creating a barrier for extrapolations of coral bleaching experiments to the real world. However, the most striking comparison was that median heating rates used in the laboratory were more than 7 times higher than those of MHWs. This suggests laboratory-based coral thermal stress responses that have been so well characterized for the past century may often represent heat shock rather than coral bleaching as it occurs in response to real-world MHWs. The accumulated heating metrics commonly used to assess heat stress on corals (i.e., DHD) may therefore be inappropriate for contrasting laboratory experiments and field observations, as temperature series with different shapes can result in the same DHDs. A short, duration experiment with a high heating rate can produce a similar DHD to a real-world MHW but represent a vastly different experience that triggers different physiological responses. In fact, different patterns in metabolic thermal performance (Martell and Zimmerman, 2021) and gene expression (Bay and Palumbi, 2015) have been observed in response to different, but rapid, heating rates in corals. Given the recent increases in duration, more so than magnitude (i.e., maximum temperature), that are driving more intense MHWs of the present day (Li and Donner, 2022; Muñiz-Castillo et al., 2019), longer, more gradual laboratory exposures are critical to predicting the coral response to climate warming.

### Measuring the Coral Bleaching Response to Heat Stress

First, most of the studies in our database measured *F*_V_/*F*_M_, indicating this response variable is heavily used in the experimental literature for coral bleaching, and in many cases, is the only variable measured as a proxy for coral bleaching. Coral bleaching is defined as in the loss of Symbiodiniaceae and/or pigments from its tissues (Brown, 1997; Douglas, 2003; Glynn, 1984; Glynn, 1993; Glynn, 1996). The most direct way to quantify coral bleaching is by isolating the tissue fraction of a piece of coral, enumerating Symbiodiniaceae cells and/or Chlorophyll *a*, and then normalizing this quantity to the coral’s surface area. All studies in our database reported air brushing or water piking methods to remove the tissue from the skeleton, with one exception. Study S00093 quantified bleaching response variables using a unique methodology that did not employ the same mechanical tissue slurrying process (Rodríguez-Troncoso et al., 2016). The study reported Symbiodiniaceae and Chlorophyll *a* values that were consistently an order of magnitude higher compared to other observations of the same species, *Pocillipora verrucosa*, and congeners from other geographies. Future studies should be aimed at comparing yields of cell counts, as the method employed by Rodríguez-Troncoso et al. (2016) may be superior to other methods (e.g., air brushing and water picking).

In contrast to these other measurements, *F*_V_/*F*_M_ does not measure the loss of Symbiodiniaceae and/or pigments, and hence is not a direct measurement of bleaching. *F*_V_/*F*_M_ is a normalized ratio achieved by dividing the variable fluorescence by the maximum fluorescence, to represent the maximum potential quantum efficiency of Photosystem II of the algae (and not a measurement of the coral animal), if all capable reaction centers are open (Baker, 2008 and references therein). The measurement is only achieved by measuring the photosynthetic subject after a period of darkness (commonly called ‘dark adaptation’ or ‘dark adapting’) to accurately interrogate the specimen’s “capacity” to capture photons. Measurements of *F*_V_/*F*_M_ are appropriate for studies concerned with changes in photobiology of photosynthesizing organisms like Symbiodiniaceae in response to changes in the environment (Brading et al., 2011; Brown et al., 1999; Emerson and Arnold, 1932; Torres et al., 2007; Warner et al., 1996; Warner et al., 2002), but less so for characterising the bleaching response in the coral holobiont to heat stress.

The frequency of measuring *F*_V_/*F*_M_ as the sole response variable in this study raises concerns that experimental studies could be mischaracterizing bleaching sensitivity to temperature and thermal stress. The widespread use of *F*_V_/*F*_M_ may be related to the convenience and accessibility of the instrumentation. In comparison with the tedious processes of cell counting and pigment extraction involved in direct bleaching measurements, *F*_V_/*F*_M_ measurements are very easy to make.

A further concern is that, despite the relative ease in collecting *F*_V_/*F*_M_ measurements, knowledge and experience is necessary to accurately use and interpret the results. First, we initially found 21 different names in our literature database used to describe the measurement that was abbreviated as *F*_V_/*F*_M_. While we removed studies that we could confirm did not measure maximum dark adapted quantum efficiency of Photosystem II from the analyses presented here, it is possible some of these descriptions were incorrectly measured or described (*sensu* Baker 2008). Second, some studies did not report whether or for how long corals were dark adapted prior to making the measurement, making those measurements invalid as well. Third, since Symbiodiniaceae have a well characterized diurnal cycle (Jones and Hoegh-Guldberg, 2001) and circadian rhythm (Sorek et al., 2013; Sorek et al., 2014), the time-of-day when *F*_V_/*F*_M_ measurements are made makes cross study and even within study comparability fraught. For example, the average maximum *F*_V_/*F*_M_ value for a healthy terrestrial leaf is ∼ 0.78 – 0.89, and for Symbiodiniaceae *in vitro* and *in hospite*, healthy values range between ∼0.58 – 0.62 (H. Martell, *pers. obs.*). These values are also strongly influenced by the location of the measurement in relation to the light incident on the coral colony (Warner and Berry-Lowe, 2006), the gain settings of the fluorometer instrument being utilized to make the measurement, and the distance of the probe from the surface of the coral. *F*_V_/*F*_M_ values greater than 0.68 in Symbiodiniaceae should be scrutinized (Ahn and Glibert, 2022; Franklin et al., 2004; Warner et al., 1996; Warner et al., 1999). In sum, our results suggest that *F*_V_/*F*_M_ may be heavily misused as a proxy for coral bleaching in the context of thermal stress and other stressors.

### Interrogating the Literature to Predict Warming Responses

Using hormesis as a case study to explore the exposures in the literature, we found the differences between laboratory and real-world exposures may substantially alter the predicted responses to warming with the use of unrealistic exposures. This raises questions about the extrapolation of experimental data in the coral conservation specifically and for all ecological studies aimed at predicting organismal survival in the context of climate change. We sought to capitalize on a broad experimental literature to detect hormesis of coral bleaching, made efforts to control for the conditions of each experimental observation in our database (i.e., *a priori* filtering the database), and found some limited evidence for coral bleaching hormesis. Although all the analyses were limited by data availability (see below), we did detect hormesis of *F*_V_/*F*_M_ when unrealistic exposures were filtered from the data via curve fitting, suggesting that hormesis may occur in the presence of realistic conditions. While the lack of observations from laboratory experiments using ecologically relevant exposures prevented us from explicitly determining whether hormesis exists, or its extent, in nature, the fact that some relationships became hormetic when unrealistic exposures were excluded from the analysis demonstrates the importance of experimental design in reducing prediction error for experiments, and for accurately predicting organismal response to climate change. It is possible hormesis requires specific conditions, such as moderate temperatures, under relatively short durations and low degree heating times, compared to the range of heating metrics in our dataset (the entire spread of the data). More likely, the lack of measurements across the range of all heating metrics influenced our results.

### Data Availability

Closing the gap between experimental and real-world in situ measurements of coral response to heat stress will require greater availability of experimental data. Only 6 % of coral bleaching studies in our database had data that were publicly available, and just 12 % of authors contacted provided temperature data, either because it was not recorded, no longer available, they were uninterested in participating, or did not respond. This drastically reduced the number of studies that could be included in analyses from 359 to 69.

The small fraction of studies with available data is the key limitation of this study and an indicator of a broad need to changes data recording and sharing practices. Some studies reported or made available response variable data without the corresponding temperature data, however no studies provided temperature data without the corresponding response variable data. This suggests a bias in author reporting, wherein response variable data are reported, but temperature data are not recorded and/or reported. More recently published studies are more likely to make data publicly available, thanks to the open science movement and expanding data availability requirements of journals. We support this openness, as well as the importance of sharing all data, especially experimental design details and input data, such as temperature.

Our comparison of heating metrics derived from temperature measurements at different scales may have introduced some error. MHW temperatures were derived from SST measurements of the “skin” of the ocean across 4 km grid cells and though SSTs are coarser spatial and temporal scales than in situ and most laboratory measurements, one study in the eastern tropical Pacific found agreement between SSTs and in situ temperatures (Claar et al., 2019), noting some bias from SSTs in coral reef grid cells toward warmer temperatures during periods of subsurface cooling and indices developed to predict coral bleaching from these data have been shown to be effective (Heron et al., 2016). Global oceanic in situ temperature datasets and comparisons don’t yet exist for coral reefs, therefore a more comprehensive dataset of such measurements would improve our estimation of MHWs on coral reefs and analyses like those presented here.

## Conclusions

Long-term laboratory experiments on corals and other marine organisms are costly and difficult to execute. The findings of this study suggest that stark differences between thermal exposures in the lab and real-world MHWs could lead to findings that are not predictive of real-world organismal responses. We encourage caution, for example, in use of short-term heat shock assays which may not be predictive of natural MHW exposures. Like others (Klein et al., 2022; McLachlan et al., 2020), we recommend longer-term experiments using ecologically relevant thermal exposures whenever possible to estimate coral bleaching thresholds and thermal performance/tolerance more accurately.

The attempt to detect hormesis in coral holobiont response to thermal exposure using a systematic review of the experimental coral bleaching literature found issues with measurement techniques, experimental design, and data availability that limit the applicability of the literature to understanding real-world coral bleaching conditions. The removal of exposures that were unrealistic substantially changed the coral physiological response to warming in almost all cases.

Based on this work, we recommend the continuous monitoring of temperature data during heating exposures and that all data be made publicly available to increase the potential for that data to be used. We also urge caution to researchers using *F*_V_/*F*_M_ as the sole measurement for coral bleaching with temperature exposure.

## DATA AVAILABILITY STATEMENT

All data, including the unfiltered and filtered databases, all protocols and scripts used for analysis in this study are available at https://github.com/harmonymartell/Temperature-Coral-Bleaching-Database. This review has not been registered. A PRISMA 2020 Checklist has been submitted with this manuscript.

## ACKNOWLEDGEMENTS

The authors thank K. Bernaus, J. Mayer, S. Mohammed, and A.U for their assistance in assembling the database and X. Li for providing the MHW data.

## FUNDING ACKNOWLEDGEMENT

This research was supported by a NSERC Discovery Grant to S. D. Donner and the NSERC CREATE Training our Future Ocean Leaders program.

